## Supplementary material for "Selection and hybridization shaped the rapid spread of African honey bee ancestry in the Americas": S2 Table - Cline model comparison

**S2 Table. Cline model comparison.** Model rankings between logistic cline fits for genome-wide *scutellata* (A) ancestry predicted by climate and distance variables.

| | predictor | df.residual | deviance | dAIC | weight |
|---|---|---|---|---|---|
| 1 | Latitude | 311 | 2.69 | 0.00 | 1.00 |
| 2 | Mean temperature | 311 | 3.63 | 93.60 | 0.00 |
| 3 | Mean temperature of coldest quarter | 311 | 5.25 | 208.60 | 0.00 |
| 4 | Minimum temperature of coldest month | 311 | 7.04 | 300.70 | 0.00 |
| 5 | Distance to Sao Paulo | 311 | 9.55 | 396.10 | 0.00 |
| 6 | Annual precipitation | 311 | 15.21 | 541.80 | 0.00 |
